# Elucidating the Impact of *Hypericum alpestre* Extract and L-NAME on the PI3K/Akt Signaling Pathway in A549 Lung Adenocarcinoma Cells

**DOI:** 10.1101/2024.05.02.592167

**Authors:** Hayarpi Javrushyan, Mikayel Ginovyan, Gayane Petrosyan, Mery Qocharyan, Tigran Harutyunyan, Smbat Gevorgyan, Zaruhi Karabekian, Alina Maloyan, Nikolay Avtandilyan

## Abstract

Plants within the *Hypericaceae* family have been traditionally used for their medicinal properties, showcasing a wide range of effects such as antibacterial, antiviral, and antioxidant qualities. *Hypericum alpestre* (HA) extracts have exhibited significant cytotoxicity against various cancer cell lines. The phenolic compounds found in HA extracts have attracted attention for their potential in cancer prevention. L-NAME, known for its ability to inhibit nitric oxide synthase (NOS) activity, has emerged as a promising approach in cancer therapy. However, the precise molecular mechanisms underlying the anticancer effects of HA and L-NAME remain unclear. This study aims to clarify the impact of HA and L-NAME on the PI3K/Akt signaling pathway in A549 lung adenocarcinoma cells, with a specific focus on TNFa/COX-2 and VEGFa/MMP-2 pathways. *In silico* analysis, they identified the compounds with the highest affinity for PI3K/Akt, a finding validated by subsequent *in vitro* experiments. Furthermore, the combination of herbs and L-NAME exhibited superior efficacy compared to the herb and 5-FU combination, as evidenced by the promotion of apoptosis. Both the herb alone and the combination of the herb with L-NAME demonstrated inhibitory effects on the TNFa/COX-2 and VEGFa/MMP-2 pathways. This therapeutic approach is hypothesized to operate through the PI3k/Akt cell signaling pathway. A better understanding of the interaction between HA polyphenols and PI3K/Akt signaling could pave the way for novel therapeutic strategies against cancer, including drug-resistant tumors.

## 1. Introduction

Hypericaceae family’s plant extracts are utilized for a variety of medical purposes, including treating wounds, bruises, dysenteries, acute mastitis, jaundices, hepatitis, sore furuncles, skin inflammation, nerve pain hemoptysis, epistaxis, metrorrhagia, irregular menstruations, burns, hemorrhages, and treatment of various tumors. The antibacterial, antiviral, antibiotic-modulating, and antioxidant/prooxidant properties of the different extracts of this plant were reported [1–5]. In our earlier studies, upon screening of ethanolic extracts of 11 wild plant species for their cytotoxic properties, *H. alpestre* (HA) extract showed one of the most potent in inhibiting the growth of A549 (lung adenocarcinoma) and HeLa (cervical carcinoma) cancer cells [5]. Phenolic compounds, as the most prevalent plant secondary metabolites, have attracted considerable attention due to their antioxidant/prooxidant properties and their potential role in the mitigation of diverse diseases associated with oxidative stress, including cancer. A total of 244 constituents were identified in *H. alpestre* extracts in our earlier research [6]. The metabolomic analysis of HA aerial part extracts using the UHPLC-ORBITRAP-HRMS technique unveiled the presence of several major phenolic constituents, suggesting that these substances might contribute to their high cytotoxic effects [7]. Considering the strong *in vitro* growth inhibiting properties of HA extract against different cancerous cell lines including the breast cancer model [7,8]. In the fight against cancer, there is a constant effort to discover new compounds with specific and targeted effects and as few side effects as possible. Currently, there is significant interest in the development of combined cancer treatments using natural compounds with chemotherapeutic agents for their modulating, drug resistance modifying properties, or reduction of chemotherapy side effects [9]. Phytochemicals can affect cellular processes and signaling pathways, which have potential antitumor properties [9].

L-NAME is an inhibitor of nitric oxide synthase (NOS), a key enzyme in several biological pathways [10]. It belongs to a group of compounds that modulate enzyme activity. L-arginine is the primary substrate for enzymes like arginase and NOS, which are linked to cancer progression [11]. This relationship underscores the potential impact of substances like L-NAME on cancer-related biological processes, highlighting the complex interplay between amino acids and enzyme function in the context of cancer [12]. Increased NOS expression is associated with different cancers such as cervical, breast, lung, brain, and spinal cord cancers [13]. Inhibition of NOS activity has been suggested as a potential tool to prevent breast cancer [14]. In our previous research, we showed that *in vivo* inhibition of NOS activity by L-NAME influences the L-arginine pathway, specifically nitric oxide and polyamine biosynthesis, which have an inhibiting effect on cancer progression [7,13,15]. For the first time, we performed a combined treatment of breast cancer with HA herb extract and L-NAME in an *in vivo* experimental model of rat breast cancer [6]. This combination decreased tumor multiplicity, increased the amount of IL-2 in the tumor environment, activated antioxidant enzymes, and decreased histological score and tumor blood vessel area. The molecular mechanisms and key cellular players, whose modification manifests such an anticancer effect, have not been clarified. In this work, we address the effects of *H. alpestre* and L-NAME on the phosphoinositide 3-kinase (PI3K)/Akt-mediated changes in A549 lung adenocarcinoma cell cultures by elucidating the roles of the TNFα/COX-2 and VEGFα/MMP-2 tandems.

The PI3K/Akt signaling pathway is a major signaling pathway in various types of cancer [16,17]. It controls the hallmarks of cancer, including cell survival, metastasis, and metabolism. The PI3K/Akt pathway also plays essential roles in the tumor environment, functioning in angiogenesis and inflammatory factor recruitment. The study of PI3K/Akt networks has led to the discovery of inhibitors for one or more nodes in the network, and the discovery of effective inhibitors is important for improving the survival of patients with cancer. To date, many inhibitors of the PI3K/Akt signaling pathways have been developed, some of which have been approved for the treatment of patients with cancer in the clinic [18,19]. However, many issues associated with the use of pathway inhibitors, including which drugs should be used to treat specific types of cancer and whether combination therapies will improve treatment outcomes, remain to be resolved. Upstream activation of the PI3K/Akt signaling pathway is essential for its function in cancer and other related diseases. Activation of this pathway is related to many factors, including TNFa and VEGFa [20]. After activation by VEGF, Akt promotes the proliferation, migration, and survival of endothelial cells, thus affecting angiogenesis. A study showed that the PI3K/Akt signaling pathway promotes the development of inflammation by affecting neutrophils, lymphocytes, and other white blood cells [21]. IL-1, IL-6, TFN-α, and other inflammatory factors activate Akt and expand the range of inflammation, while Akt inhibition blocks both inflammation and tumor development [17]. Akt signaling promotes tumor cell survival, proliferation, growth, and metabolism by activating its downstream effectors such as COX-2 MMP-2, and Caspase-3. Multiple signaling pathways are involved in the regulation of tumor angiogenesis, among which PI3K/Akt signaling is the most important. PI3K forms a complex with E-cadherin, β-catenin, and VEGFR-2 and is involved in endothelial signaling mediated by VEGF through the activation of the PI3K/Akt pathway[22]. The PI3K/Akt signaling pathway also promotes TNF-induced endothelial cell migration and regulates tumor angiogenesis. Matrix metalloproteinases (MMPs) and cyclooxygenase 2 (COX-2) also affect tumor angiogenesis [21,23]. In tumor invasion and metastasis, platelet-derived growth factor (PDGF) induces MMP expression through a PI3K-mediated signaling pathway. Upregulation of the antiapoptotic protein Bcl-2 and activation of the PI3K/Akt signaling pathway are the main mechanisms by which COX-2 stimulates endothelial angiogenesis. PI3K/Akt signaling blocks the expression of proapoptotic proteins, reduces tissue apoptosis, and increases the survival rate of cancer cells. Akt inhibits the proapoptotic factors Bad and procaspase-9 through phosphorylation and inhibits Caspas-3 activity [16,17].

Based on the above facts, we have set ourselves the task of elucidating, through *in silico* methods, the polyphenols present in the herb that exhibit the highest affinity for PI3K/Akt. To ensure the reliability of our data, we used *in vitro* A549 lung adenocarcinoma cell cultures to demonstrate the herb extract’s effects on PI3K and Akt enzymes and clarify how PI3K/Akt mediation regulates TNFα, VEGFα, COX-2, MMP-2, and Caspase-3 components. Since PI3K/Akt signaling dysregulation plays a crucial role in cancer drug resistance, discovering new compounds targeting these components could help overcome drug resistance in various cancer therapies.

## 2. Materials and Methods

### 2.1. Chemicals and reagents

All chemicals were purchased from Sigma-Aldrich (USA) and Abcam (UK). Antibodies against TNFa (ab46087), VEGFa (ab193555), MMP-2 (ab92536), COX-2 (ab38898), PI3K and phosphorylated (p)-PI3K (ab191606), as well as ELISA kits for AKT and p-AKT (ab179463) were purchased from Abcam.

### 2.2. Plant material

The aerial parts of the Hypericum alpestre plant were harvested from the Tavush region of Armenia (at an altitude of 1600-2800 meters above mean sea level) during the flowering period, following the previously described protocol [24]. Identification of plant materials was carried out at the YSU Department of Botany and Mycology by Dr. Narine Zakaryan. The plant materials were deposited at the Herbarium of Yerevan State University (YSU), where they were assigned a voucher specimen serial number [7].

### 2.3. Plant crude extract preparation

The grounded plant materials were extracted by maceration with 96% ethanol at a 10:1 solvent-to-sample ratio (v/w). Stock solutions of 50 mg DW/mL crude ethanol extract were prepared as described earlier [25]. The percent yield was 10.60 ± 2.31%.

### 2.4. Cell cultures

Human lung adenocarcinoma A549 cell culture was obtained from ATCC (cat # CCL-185) and maintained in DMEM medium supplemented with L-glutamine (2 mmol/L), sodium pyruvate (200 mg/L), fetal bovine serum (100 mL/L), and antibiotics (100 U/mL penicillin and 100 µg/L streptomycin). Cells were grown at 37 °C under a humidified atmosphere with 5% CO_2_ in a Biosmart (Biosan, Latvia) as described before [26]. Cultured cells were regularly examined for the presence of mycoplasma contamination using the Universal Mycoplasma Detection Kit from ATCC (Manassas, Virginia, USA).

### 2.5. MTT cytotoxicity test

The MTT test was performed as described previously [27] to assess the growth inhibition of A549 cells exposed to different concentrations of the *H. alpestre* extract for 4, 24, or 72 h.

### 2.6. ELISA of TNFa, VEGFa, COX-2, MMP-2, and Akt

A549 cells (2 × 10^5^) were cultured in 12-well plates and incubated for 24 h. After incubation, the cell medium (630 *μ*L) was replaced and the cells were treated with PBS and 1% Ethanol solution (Control, A549), 5-FU (40*μ*M), HA (0.25mg/mL), L-NAME (14mM, LN) and HA+L-NAME (0.25mg/mL + 14mM, HALN for 24 h and then the culture medium was harvested. TNFa, VEGFa, and MMP-2 in the supernatant were quantified according to the manufacturer’s instructions. Cells from each group were collected (trypsinized, neutralized, centrifuged), lysed on ice with Lysis buffer, collected in a centrifuge tube, and further lysed for 10 min. The supernatant was collected after centrifugation at 13,000 × g for 10 min at 4°C. Changes in the levels of COX-2 and Akt were measured using ELISA kits, according to the manufacturer’s instructions. Protein concentration in cell culture medium and lysates were measured with a Bradford method. Each test sample (70 µL) was added from three biologically independent repetitions, involving three different cell passages, with each passage containing two technical replicates.

### 2.7. Phospho-PI 3 kinase p85 + Total In-Cell ELISA assay

A549 cells (1.5×10^4^ cells per well) were seeded in the 96-well plates treated for tissue culture. After 24 h incubation, the cell medium (180 *μ*L) was refreshed. The cells were treated with 20*μ*L control or test compounds with the following final concentrations: PBS, 1% ethanol solution (Control, A549C), 5-FU (40*μ*M), HA (0.25mg/mL), L-NAME (14mM, LN) and HA+L-NAME (0.25mg/mL + 14mM, HALN). The calculations for seeding the cells were conducted to ensure approximately 80% confluency at the time of fixation. After 24 h exposure, the medium was discarded and cells were fixed with 100 µL of 4% formaldehyde in PBS. Crystal Violet was used to stain cells for normalizing readings in 450nm for Phospho-PI 3 kinase p85 + Total. The measured OD450 readings were normalized for cell number by dividing each OD450 reading by the corresponding OD595 reading for the same well. This relative cell number was then used to normalize each reading. Total and phospho-PI 3 kinase p85 were each assayed in triplicate using the phospho- and total PI 3 Kinase p85 antibodies included in the PI 3 Kinase Kit. Levels of Phospho-PI 3 kinase p85 and Total PI3K were measured using an In-Cell ELISA kit (ab207484), according to the manufacturer’s instructions.

### 2.8. Caspase-3/CPP32 Colorimetric assay

A549 cells (5 × 10^5^ cells per well) were cultured in 6-well plates and incubated for 24 h. Then, the cell medium (900 *μ*L) was refreshed and the cells were treated with 10 *μ*L of PBS+1 % ethanol solution (control, A549C) or test compounds with the following final concentrations: 5-FU (40 *μ*M), HA (0.25 mg/mL), L-NAME (14mM, LN), HA+L-NAME (0.25mg/mL + 14mM, HALN) and HA+5-FU (0.25 mg/mL + 40 *μ*M). After 24 h the cells were harvested. Each test sample (100 µL) was added from three biologically independent repetitions, involving three different cell passages, with each passage containing two technical replicates. Cells were resuspended in 50 µL of chilled Cell Lysis Buffer and incubated on ice for 10 minutes. Then, cell lysate was centrifuged for 1 min (10,000 x g). After that supernatant (cytosolic extract) was transferred to a fresh tube and put on ice for immediate assay. Fold-increase in CPP32 activity has been determined by comparing these results with the level of the uninduced control. Optical density values were corrected taking into account the number of cells. All steps were performed according to the protocol presented in the Caspase-3/CPP32 Colorimetric Assay Kit (K106, BioVision) instructions.

### 2.9. Morphological analysis of apoptosis by Hoechst 33258 staining

Hoechst 33258 staining enables the determination of the percentage of apoptotic cells through morphological analysis using a fluorescence microscope [28]. Briefly, A549 cells (2×10^5^ cells/mL) were incubated with vehicle or 5-FU (40 *μ*M) as a positive control, and test samples: HA (0.25 mg/mL), LN (14 mM), HA+5-FU (0.25 mg/mL + 40 *μ*M), and HA+LN (0.25mg/mL+14mM) for 24 h. Cells were fixed with 4 % paraformaldehyde in PBS for 10 min, washed twice with PBS for 5 min, and stained with Hoechst 33258 (10 *μ*g/mL) for 10 min in the dark. Then cells were analyzed under a fluorescence microscope (x250 magnification) (Zeiss, Germany). Cells with typical morphological changes of nuclei, such as chromatin condensation, rough edges, nuclear fragmentation, and apoptotic bodies were counted as apoptotic. Controls and treatment variants were examined in duplicate. For each variant, 500 cells were scored and the percentage of apoptotic cells was calculated as follows: % apoptotic cells = (the number of apoptotic cells/500 cells)*100.

### 2.10. Acquisition and Analysis of Protein Structures

The structural elucidation of PI3K and AKT protein was conducted by retrieving their crystallographic forms from the Protein Data Bank (PDB), accessed at [https://www.rcsb.org/], with the specific PDB IDs being 6AUD and 2JDO, respectively. Advanced molecular visualization and analysis were performed using the PyMOL Molecular Graphics System (Schrödinger, LLC). This phase included preprocessing steps for the removal of non-essential entities such as solvent molecules, ions, and extraneous non-protein components. Concurrently, ligands associated with these protein structures were segregated, yielding pure protein structures for subsequent computational analysis. The refined protein models were earmarked for molecular docking studies, whereas the separated ligands were preserved for re-docking procedures in subsequent structural bioinformatics applications.

### 2.11. Enhanced Molecular Docking Using AutoDock Vina

AutoDock Vina, an extensively utilized molecular docking tool, employs the Iterated Local Search global optimizer. This optimizer is conceptually akin to those used in ICM and the Broyden-Fletcher-Goldfarb-Shanno (BFGS) quasi-Newton algorithm for local optimization. The software integrates a hybrid scoring function that combines empirical and knowledge-based approaches to enhance docking precision and efficiency. For optimal docking performance, the ‘exhaustiveness’ parameter was set to 8, adhering to the standard protocols recommended by the software developers. Utilizing AutoDock Vina facilitated a comprehensive assessment of the binding affinities of key compounds extracted from Hypericum with the target proteins, allowing for their prioritization based on predicted binding efficacy [29]

### 2.12. Statistic analysis

Results are presented as means ± SEM. We analyzed the data using either one-way ANOVA or its non-parametric counterpart, the Kruskal-Wallis test, depending on the results of the normality test. Subsequently, Dunn’s test was employed to evaluate the statistical significance of the results for TNFα, VEGFα, MMP-2, COX-2, Caspase-3, and apoptosis rate. The significance of the results obtained for PI3K and Akt was assessed using two-way ANOVA and Tukey’s multiple comparisons tests. Statistical analyses were performed using GraphPad Prism 8 software (San Diego, CA, USA), and a significance level of p<0.05 was deemed statistically significant.

## 3. Results

### 3.1. The interaction of HA extract metabolites with PI3K and Akt

Our previous work elucidated more than 200 active polyphenolic compounds in HA [7]. In this study, the interaction of these compounds and PI3k/Akt was clarified by in silico methods. 2 compounds with the highest affinity were identified. Our results detail the specific binding interactions of chrysoeriol glucuronide and pseudohypericin with PI3K and AKT proteins, as observed in our molecular docking simulations (Fig. 1, A, B, C, and D). For chrysoeriol glucuronide docked with PI3K, multiple hydrogen bonds were identified: the ligand formed a hydrogen bond with the backbone nitrogen of Ala805 at a distance of 3.14 Å, and additional hydrogen bonds with Ser806 and Thr886, indicative of a strong interaction potential (Fig. 1, B). The ligand also exhibited hydrophobic interactions with key residues including Trp812 and Met804, which are likely to contribute to the binding affinity and specificity. Docking chrysoeriol glucuronide with AKT resulted in a distinct interaction profile, with fewer hydrogen bonds observed (Fig. 1, A). Notably, a hydrogen bond with Glu230 at 2.70 Å was prominent, alongside hydrophobic contacts with Lys181 and Val166, suggesting an alternate binding conformation compared to its binding with PI3K. Pseudohypericin docking with PI3K revealed a different pattern: a network of hydrogen bonds with Glu880, Asp964, and Ile831, and hydrophobic interactions with residues such as Val882 and Tyr867 (Fig. 1, D). This pattern indicates a binding mode that is distinct from that of chrysoeriol glucuronide with PI3K. Finally, pseudohypericin docked with AKT showed an interaction pattern that includes hydrogen bonds with Asp293 and Lys160, and hydrophobic contacts with Glu236 and Met282 (Fig. 1, C). The interaction profile was again unique from its binding with PI3K and chrysoeriol glucuronide’s interactions with both kinases.

**Fig. 1.**
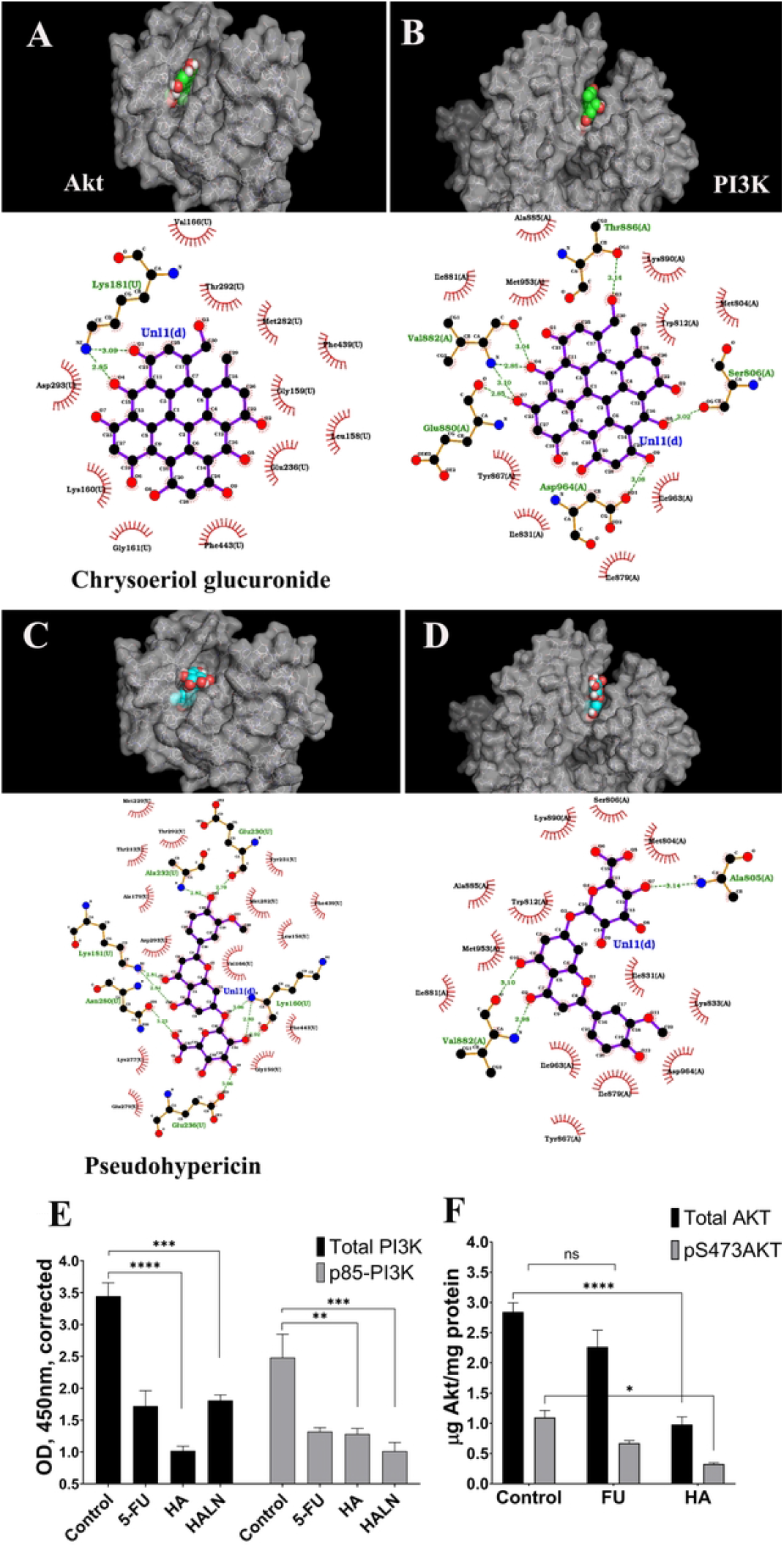
2D binding analysis and interaction types on PI3K (B, D) and AKT (A, C) in combination with 3D visualization. A and B – Chrysoeriol glucuronide, C and D – Pseudohypericin. Hydrogen bonding is indicated by green dotted lines, while the remaining interactions are hydrophobic. Effect of HA, 5-FU, L-NAME, HA+L-NAME on PI3K/Akt pathway in A549 cells (E - PI3K, F - Akt). Total and phospho-kinases were each assayed in triplicate using the phospho- and total Kinase antibodies included in the PI 3 Kinase and Akt kits (n=3, * - p≤0.05, ** - p≤0.01, *** - p≤0.001, **** - p≤0.0001). p85-PI3K - Phospho-PI 3 kinase p85, pS473AKT - phospho-Akt (Ser473).

These results highlight the specificity of ligand-kinase interactions and suggest that chrysoeriol glucuronide and pseudohypericin could potentially act as selective kinase inhibitors due to their ability to form multiple hydrogen bonds and hydrophobic interactions with distinct residues in PI3K and AKT (Table 1 and 2).

**Table 1.** Docking results of HA major bioactive constituents on PI3K and Akt.

| **Ligand** | **PI3K**<br>**Binding_Affinity** | **AKT**<br>**Binding_Affinity** | **Total** |
| --- | --- | --- | --- |
| **Pseudohypericin** | -12.7 | -7.6 | -20.3 |
| **Chrysoeriol-glucuronide** | -8.7 | -9.7 | -18.4 |
| **Trihydroxyxanthone** | -8.6 | -8.6 | -17.2 |
| **Biapigenin** | -8 | -8.9 | -16.9 |
| **Quercetin** | -8.4 | -8.5 | -16.9 |
| **Myricetin_glucoside** | -8.5 | -8.3 | -16.8 |
| **Myricetin** | -8.7 | -8 | -16.7 |
| **References** | -9.6 | -7 | -16.6 |
| **Quercetin-acetyl-glucoside** | -8.5 | -8 | -16.5 |
| **Gallocatechin** | -8.3 | -8 | -16.3 |
| **Catechin** | -8.5 | -7.8 | -16.3 |
| **Quercetin-pentoside_2D_3D** | -8.5 | -7.8 | -16.3 |
| **Dihydromyricetin** | -8.1 | -8.1 | -16.2 |
| **Myricetin-arabinoside** | -8.5 | -7.4 | -15.9 |
| **Caffeoylquinic_acid** | -7.8 | -8.1 | -15.9 |
| **Epicatechin** | -7.8 | -8.1 | -15.9 |
| **p-Coumaroylquinic_acid** | -7.7 | -7.7 | -15.4 |
| **Galloyl_glucose** | -7.7 | -7.6 | -15.3 |
| **Quercetin-hexoside_2D_3D** | -7.5 | -7.7 | -15.2 |
| **Oleuropein** | -7.9 | -7.3 | -15.2 |
| **Vanillic_acid_glucoside** | -7.3 | -7.5 | -14.8 |
| **Carnosic_acid** | -7.5 | -7.2 | -14.7 |
| **Salicylic_acid_glucoside** | -7.2 | -7.4 | -14.6 |
| **Hypercalin_A** | -7.5 | -7.1 | -14.6 |
| **Methoxy-carnosic_acid** | -7.4 | -6.9 | -14.3 |
| **Hyperevolutin_A** | -6.1 | -7.5 | -13.6 |
| **Procyanidin_dimer** | -6.3 | -7.1 | -13.4 |
| **Trihydroxy_octadecadienoic_acid** | -6.6 | -6.5 | -13.1 |
| **Chipericumin_C** | -5.6 | -6.7 | -12.3 |
| **Pinellic_acid** | -6.2 | -6.1 | -12.3 |
| **hyperforin** | -4.4 | -7 | -11.4 |
| **Hyperpapuanone** | -6.1 | -5.2 | -11.3 |
| **adhyperforin** | -4.2 | -6.8 | -11 |
| **Quinic_acid** | -5.2 | -5.2 | -10.4 |
| **skyrin_glucoside_3D** | -6.9 | -3.5 | -10.4 |
| **Gluconic_acid** | -4.2 | -4.7 | -8.9 |

**Table 2.**
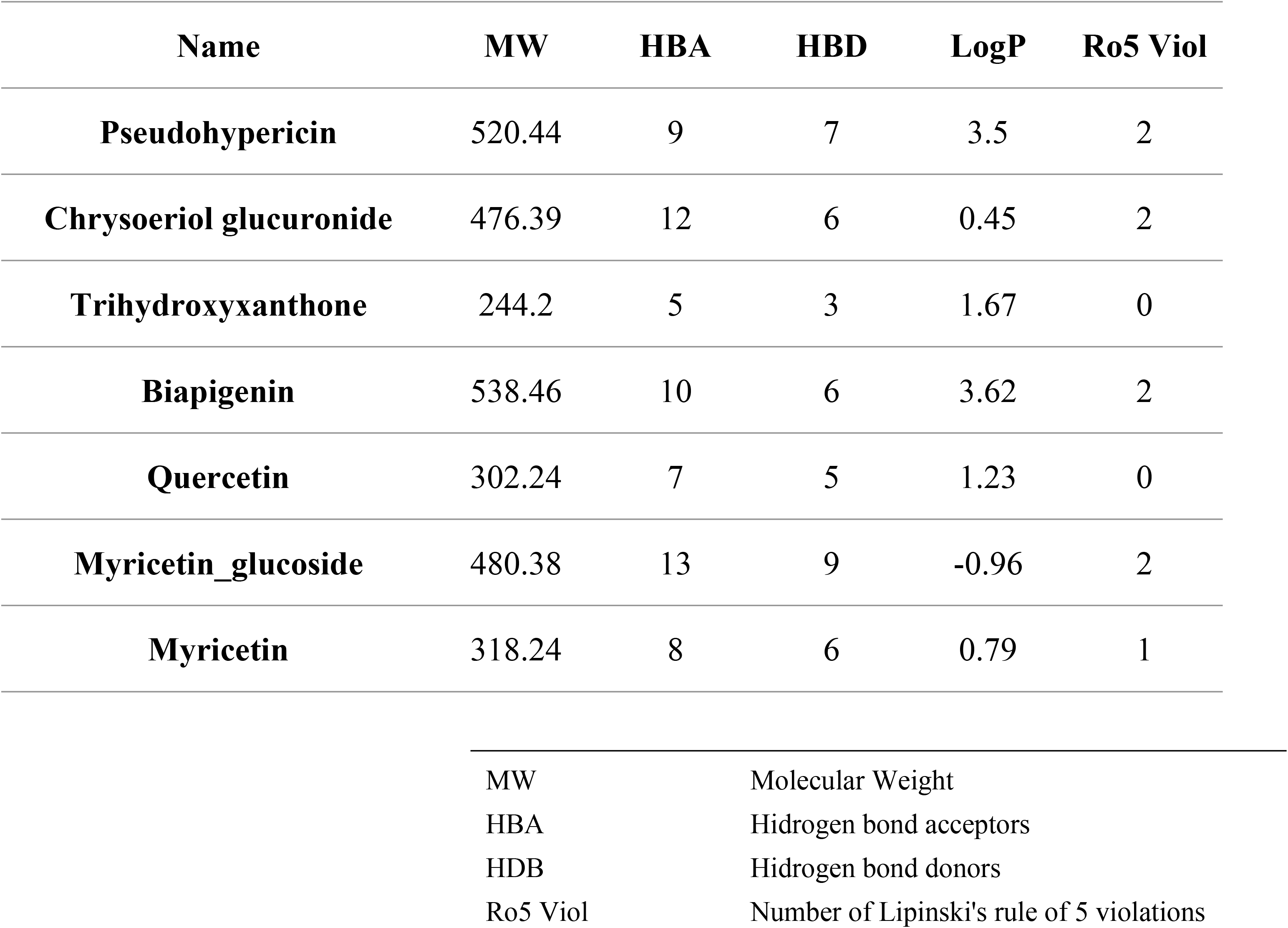
ADME properties of HA extract major phytochemicals.

| Name | MW | HBA | HBD | LogP | Ro5 Viol |
| --- | --- | --- | --- | --- | --- |
| Pseudohypericin | 520.44 | 9 | 7 | 3.5 | 2 |
| Chrysoeriol glucuronide | 476.39 | 12 | 6 | 0.45 | 2 |
| Trihydroxyxanthone | 244.2 | 5 | 3 | 1.67 | 0 |
| Biapigenin | 538.46 | 10 | 6 | 3.62 | 2 |
| Quercetin | 302.24 | 7 | 5 | 1.23 | 0 |
| Myricetin_glucoside | 480.38 | 13 | 9 | -0.96 | 2 |
| Myricetin | 318.24 | 8 | 6 | 0.79 | 1 |

|  |  |
| --- | --- |
| MW | Molecular Weight |
| HBA | Hydrogen bond acceptors |
| HDB | Hydrogen bond donors |
| Ro5 Viol | Number of Lipinski's rule of 5 violations |

In the next research phase, the possible effect of the herb extract on PI3K and Akt in A549 cancer cells was clarified *in vitro*. The results show that the extract of the herb and the combination with L-NAME reduces both the total amount of PI3K and its phosphorylated version (Fig. 1, E).

The latter shows that it can influence both the activity of the enzyme and its gene expression. This fact confirms the results obtained by the *in silico* study. The observation of Akt showed that the herb reduces both the total and phosphorylated forms of the enzyme (Fig. 1, F).

### 3.2. *H. alpestre* extracts alone and in combination with L-NAME downregulate quantitative changes of TNFa, VEGFa, COX-2, and MMP-2

To clarify the factors influencing PI3K/Akt inhibition, we observed the quantitative changes in TNFα and VEGFα. *H. alpestre* extract reduced the amount of TNFα in A549 cells by approximately two-fold (Fig. 2, A), and decreased the level of VEGFα by 2.7 times (Fig. 2, B). These two factors, being interconnected with PI3K/Akt enzymes, influence and accept the decrease in their activity. To understand which compounds are affected by the signal after Akt, COX-2 and MMP-2 were considered. HA reduced the amount of COX-2 by about 30% (Fig. 2, C), and the amount of MMP-2 by 20% (Fig. 2, D). In the case of COX-2, the combined effect of *H. alpestre* and L-NAME is synergistic, reducing its level by approximately five-fold (Fig. 2, C). The amount of MMP-2 decreased by 35% under the influence of HALN (Fig. 2, D).

**Fig. 2.**
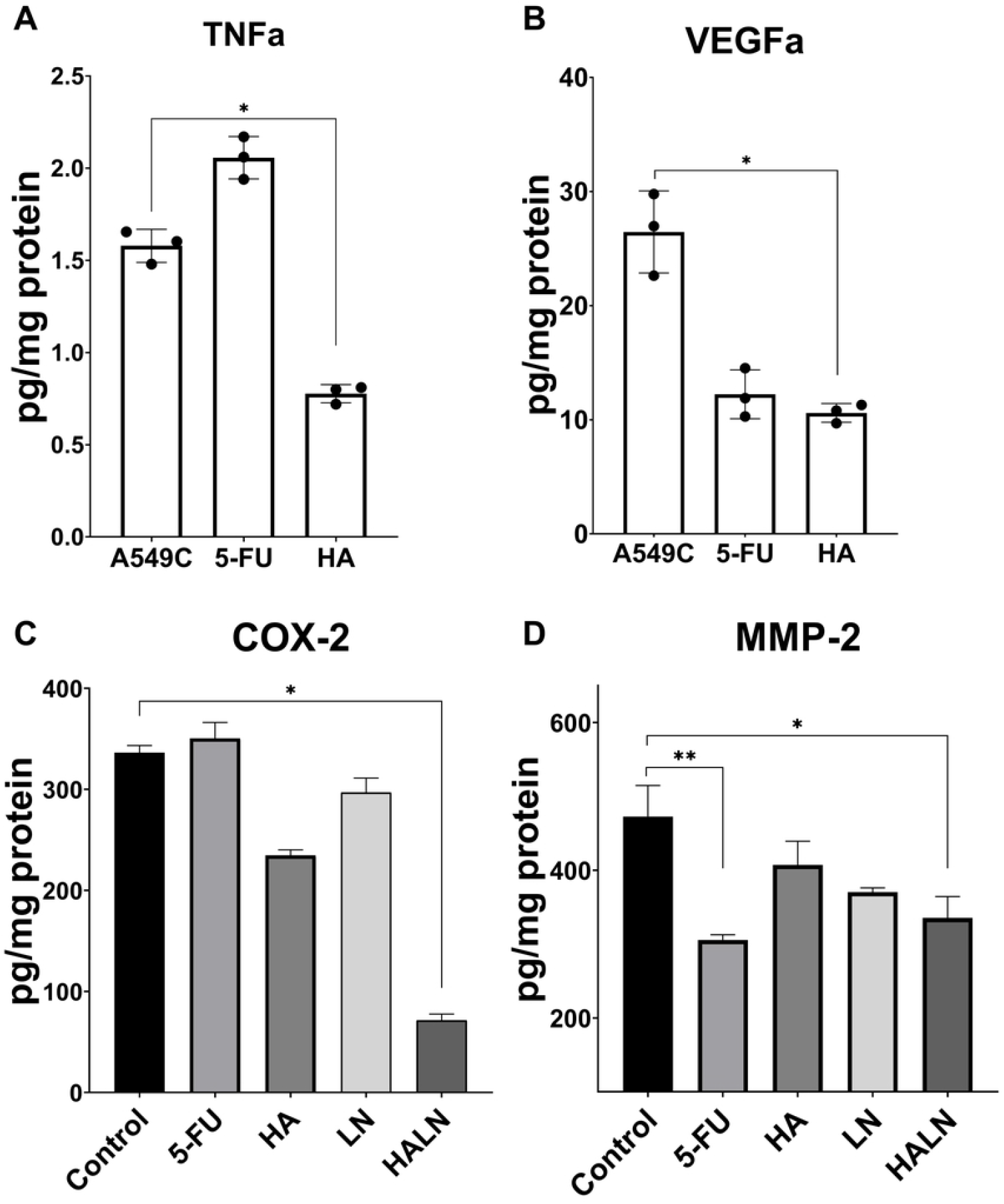
The influence of *H. alpestre* extract alone and in combination with L-NAME on quantitative changes of TNFa (A), VEGFa (B), COX-2 (C), and MMP-2 (D) in A549 cells. Control - A549C, 5-Fluorouracil - 5-FU (40*μ*M), *H. alpestre* - HA (0.25mg/mL), L-NAME – 14mM LN, HALN - HA+L-NAME (0.25mg/mL + 14mM). Three biologically independent repetitions of the experiments were performed, with two technical replicates each (n=3, * - p≤0.05, ** - p≤0.01).

### 3.3. Assessment of apoptosis

In Hoechst 33258 staining apoptotic cells can be morphologically distinguished from viable cells by karyopyknosis, rough edges of their nuclei, and signs of nuclear fragmentation (**Fig. 3, marked with arrows**).

**Fig. 3.**
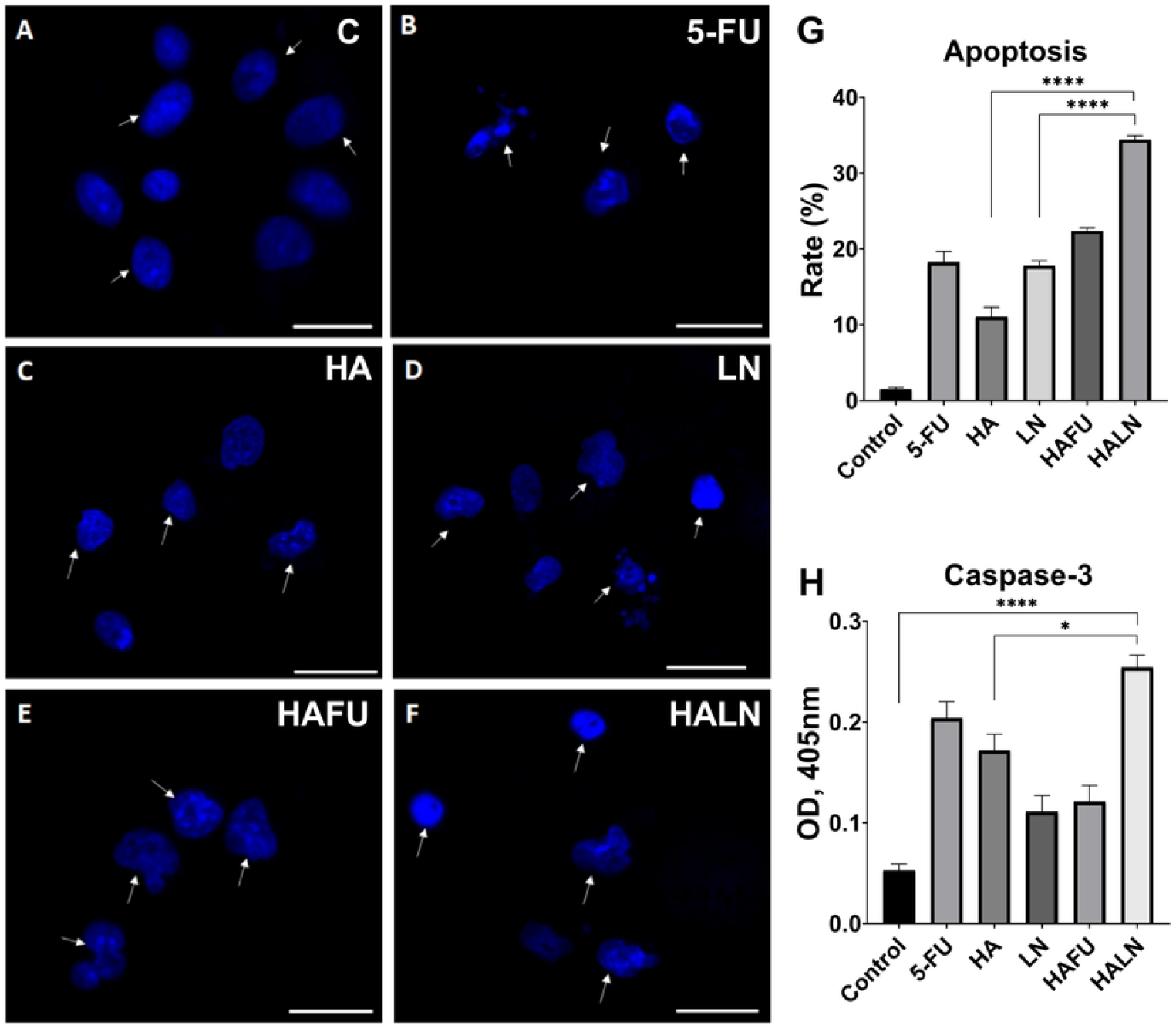
Assessment of apoptosis by Hoechst 33258 staining (blue) and caspase-3 activity in A549 cells. Arrowheads indicate apoptotic cell nuclei. The scale bar is 100 *μ*m. (A) Control cell nuclei have smooth edges. Treatment of cells with 5-FU (B), HA (C), LN (D), 5-FU+HA (E), and HA+LN (F) induced chromatin condensation, nuclear shrinkage, or fragmentation. (G) Frequencies of apoptotic cells detected by morphological analysis using Hoechst 33258 staining, ^*^p<0.05, ^**^p<0.01, ***p<0.001, ****p<0.0001.

All treatment variants induced a significant elevation of apoptosis rate compared to untreated control (**Fig. 3**) (p<0.05). In control cells, the rate of apoptosis was 2.30±1.51 %, while 24 h incubation with 5-FU increased the rate of apoptotic cells to 15.61±1.94% (**Fig. 3, B and G**). Treatments with HA or LN alone increased the rates of apoptotic cells up to 17.00±1.41% and 17.20±0.80%, respectively (**Fig. 3, C, D, and G**). Treatment of cells with combinations of 5-FU+HA or HA+LN increased the rates of apoptotic cells up to 18.90±0.70% and 27.7±0.70%, respectively (**Fig. 3, E, and F**). Thus, the apoptosis of A549 cells induced by the combination of HA+LN was significantly higher compared to that induced by 5-FU, HA, LN, and the combination of 5-FU+HA. (p<0.05) (**Fig. 3, G**). We further evaluated the expression of active (cleaved) Caspase-3 using a colorimetric assay (**Fig. 3, H**). The increase in Caspase-3 activity was observed in all treatment variants which was more pronounced in cells treated with HA+LN (p<0.05).

## 4. Discussion

The study investigates the potential synergistic effects of *Hypericum alpestre* (HA) extract and L-NAME (Nω-Nitro-L-arginine methyl ester), an inhibitor of nitric oxide synthase (NOS), on the PI3K/Akt signaling pathway in A549 lung adenocarcinoma cells. The research employed molecular docking simulations to identify potential compounds from HA extract that exhibit high affinity for PI3K and Akt proteins. Chrysoeriol glucuronide and pseudohypericin emerged as promising candidates based on their strong interactions with PI3K and Akt. These findings provide valuable insights into the molecular mechanisms underlying the anticancer effects of HA extract.

Subsequent *in vitro* experiments confirmed the inhibitory effects of HA extract on PI3K and Akt enzymes in A549 cells. Treatment with HA extract alone or in combination with L-NAME resulted in a significant reduction in both total and phosphorylated forms of PI3K and Akt. This suggests that the HA extract may exert its anticancer effects through modulation of the PI3K/Akt signaling pathway. However, other mechanisms also can be involved. The study observed quantitative changes in key signaling molecules associated with cancer progression. HA extract alone significantly reduced the levels of TNFa, VEGFa, COX-2, and MMP-2 in A549 cells. Furthermore, the combination of HA extract and L-NAME exhibited a synergistic effect in downregulating COX-2 and MMP-2 expression, indicating a potential mechanism for enhancing the anticancer properties of HA extract.

Treatment with HA extract alone or in combination with L-NAME significantly increased apoptosis in A549 cells, as evidenced by morphological changes and elevated Caspase-3 activity. The combination of HA extract and L-NAME demonstrated superior efficacy in inducing apoptosis compared to either treatment alone or conventional chemotherapy (5-FU), highlighting the potential of this combination as a novel therapeutic strategy for lung adenocarcinoma.

The study provides valuable insights into the molecular mechanisms underlying the anticancer effects of HA extract and L-NAME combination therapy. These findings support further preclinical and clinical investigations to evaluate the efficacy and safety of this combination in lung adenocarcinoma and other cancer types. Additionally, the identification of specific compounds from HA extract with high affinity for PI3K/Akt proteins may facilitate the development of targeted therapies with improved efficacy and reduced side effects.

## 5. Conclusion

The study elucidated the molecular mechanisms underlying these properties by focusing on the PI3K/Akt signaling pathway and its downstream targets, including TNFa/COX-2 and VEGFa/MMP-2 pathways. *In silico* analysis identified specific compounds from HA extract, such as chrysoeriol glucuronide and pseudohypericin, with high affinity for PI3K and Akt proteins. Subsequent *in vitro* experiments confirmed that HA extract and its combination with L-NAME reduced both the total amount and phosphorylated forms of PI3K and Akt, indicating their inhibitory effect on the PI3K/Akt pathway. Moreover, treatment with HA extract alone and in combination with L-NAME led to significant downregulation of TNFa, VEGFa, COX-2, and MMP-2, which are key components associated with cancer progression and angiogenesis. The observed increase in apoptosis, as evidenced by morphological changes and elevated Caspase-3 activity, further supports the efficacy of HA extract and L-NAME combination therapy inducing cancer cell death.

Overall, these findings suggest that the synergistic effects of HA extract and L-NAME on the PI3K/Akt signaling pathway and downstream targets hold promise for developing novel therapeutic strategies against cancer, particularly for drug-resistant tumors. Further research is warranted to validate these findings and explore the clinical potential of this combination therapy.

## 6. Availability of data and materials

The data used to support the findings of this study are included in the articles.

## 8. Authors’ contributions

The study’s conception and design were the results of collective contributions from all authors. The investigations and analysis of results were carried out by MG, NA, HJ, PG, MQ, and TH. MG and NA wrote the manuscript. Assessment of apoptosis rate by Hoechst 33258 staining and analysis of apoptosis performed by TH. The docking and ADME of the top compounds present in the HA ethanolic extract were performed by SG. NA, MG, HJ, and AM directed the project, corrected, and edited the manuscript. All authors participated in the revision and approval of the final version of the manuscript.

## 9. Acknowledgments

Plant materials were identified by Dr. Narine Zakaryan from the Department of Botany and Mycology at Yerevan State University (YSU). This work was supported by the Science Committee of MESCS RA through research projects numbered 21AG-1F068 and 23LCG-1F010.

## References

1. Ginovyan MM, Sahakyan NZ, Petrosyan MT, Trchounian AH. Antioxidant potential of some herbs represented in Armenian flora and characterization of phytochemicals. Proceedings of the YSU B: Chemical and Biological Sciences. 2021;55: 25–38.

2. Ginovyan M, Ayvazyan A, Nikoyan A, Tumanyan L, Trchounian A. Phytochemical Screening and Detection of Antibacterial Components from Crude Extracts of Some Armenian Herbs Using TLC-Bioautographic Technique. Curr Microbiol. 2020;77: 1223–1232. doi:10.1007/s00284-020-01929-0

3. Ginovyan M, Andreoletti P, Cherkaoui-Malki M, Sahakyan N. Hypericum alpestre extract affects the activity of the key antioxidant enzymes in microglial BV-2 cellular models. AIMS Biophys. 2022;9: 161–171. doi:10.3934/biophy.2022014

4. Ginovyan M, Javrushyan H, Karapetyan Hasmik, Avtandilyan Nikolay. Hypericum alpestre extract exhibits in vitro and in vivo anticancer properties by regulating the cellular antioxidant system and metabolic pathway of L-arginine. Cell Biochemistry & Function. 2023.

5. Ginovyan M, Hovhannisyan S, Javrushyan H, Sevoyan G, Karabekian Z, Zakaryan N, et al. Screening revealed the strong cytotoxic activity of Alchemilla smirnovii and Hypericum alpestre ethanol extracts on different cancer cell lines. AIMS Biophys. 2022;10: 12–22. doi:10.3934/BIOPHY.2023002

6. Ginovyan M, Javrushyan H, Karapetyan H, Koss-Mikolajczyk I, Kusznierewicz B, Grigoryan A, et al. Hypericum alpestre extract exhibits in vitro and in vivo anticancer properties by regulating the cellular antioxidant system and metabolic pathway of L-arginine. Cell Biochem Funct. 2024;42. doi:10.1002/cbf.3914

7. Ginovyan M, Javrushyan H, Karapetyan H, Koss-Mikolajczyk I, Kusznierewicz B, Grigoryan A, et al. Hypericum alpestre extract exhibits in vitro and in vivo anticancer properties by regulating the cellular antioxidant system and metabolic pathway of L-arginine. Cell Biochem Funct. 2024;42. doi:10.1002/cbf.3914

8. Ginovyan M, Hovhannisyan S, Javrushyan H, Sevoyan G, Karabekian Z, Zakaryan N, et al. Screening revealed the strong cytotoxic activity of Alchemilla smirnovii and Hypericum alpestre ethanol extracts on different cancer cell lines. AIMS Biophys. 2022;10: 12–22. doi:10.3934/BIOPHY.2023002

9. Chavda VP, Patel AB, Mistry KJ, Suthar SF, Wu Z-X, Chen Z-S, et al. Nano-Drug Delivery Systems Entrapping Natural Bioactive Compounds for Cancer: Recent Progress and Future Challenges. Front Oncol. 2022;12. doi:10.3389/fonc.2022.867655

10. Pershing NLK, Yang C-FJ, Xu M, Counter CM. Treatment with the nitric oxide synthase inhibitor L-NAME provides a survival advantage in a mouse model of Kras mutation-positive, non-small cell lung cancer. Oncotarget. 2016;7: 42385–42392. doi:10.18632/oncotarget.9874

11. Chioda M, Marigo I, Mandruzzato S, Mocellin S, Bronte V. Arginase, Nitric Oxide Synthase, and Novel Inhibitors of L-arginine Metabolism in Immune Modulation. Second Edi. Cancer Immunotherapy: Immune Suppression and Tumor Growth: Second Edition. Elsevier; 2013. doi:10.1016/B978-0-12-394296-8.00034-8

12. Szabo C. Gasotransmitters in cancer: from pathophysiology to experimental therapy. Nat Rev Drug Discov. 2016;15: 185–203. doi:10.1038/nrd.2015.1

13. Avtandilyan N, Javrushyan H, Ginovyan M, Karapetyan A, Trchounian A. Anti - cancer effect of in vivo inhibition of nitric oxide synthase in a rat model of breast cancer. Mol Cell Biochem. 2022. doi:10.1007/s11010-022-04489-y

14. Zhang L, Zeng M, Fu BM. Inhibition of endothelial nitric oxide synthase decreases breast cancer cell MDA-MB-231 adhesion to intact microvessels under physiological flows. American Journal of Physiology-Heart and Circulatory Physiology. 2016;310: H1735–H1747. doi:10.1152/ajpheart.00109.2016

15. Ginovyan M, Javrushyan H, Petrosyan G, Kusznierewicz B, Koss-Mikołajczyk I, Koziara Z, et al. Anti-cancer effect of Rumex obtusifolius in combination with arginase/nitric oxide synthase inhibitors via downregulation of oxidative stress, inflammation, and polyamine synthesis. Int J Biochem Cell Biol. 2023;158: 106396. doi:10.1016/j.biocel.2023.106396

16. Fruman DA, Chiu H, Hopkins BD, Bagrodia S, Cantley LC, Abraham RT. The PI3K Pathway in Human Disease. Cell. Cell Press; 2017. pp. 605–635. doi:10.1016/j.cell.2017.07.029

17. He Y, Sun MM, Zhang GG, Yang J, Chen KS, Xu WW, et al. Targeting PI3K/Akt signal transduction for cancer therapy. Signal Transduction and Targeted Therapy. Springer Nature; 2021. doi:10.1038/s41392-021-00828-5

18. Bo S, Lai J, Lin H, Luo X, Zeng Y, Du T. Purpurin, a anthraquinone induces ROS-mediated A549 lung cancer cell apoptosis via inhibition of PI3K/AKT and proliferation. Journal of Pharmacy and Pharmacology. 2021;73: 1101–1108. doi:10.1093/jpp/rgab056

19. Shi F, Zhang J, Liu H, Wu L, Jiang H, Wu Q, et al. The dual PI3K/mTOR inhibitor dactolisib elicits anti-tumor activity in vitro and in vivo. Oncotarget. 2018. Available: http://www.impactjournals.com/oncotarget

20. Jiang BH, Liu LZ. Chapter 2 PI3K/PTEN Signaling in Angiogenesis and Tumorigenesis. Advances in Cancer Research. 2009. pp. 19–65. doi:10.1016/S0065-230X(09)02002-8

21. Yang J, Wang X, Gao Y, Fang C, Ye F, Huang B, et al. Inhibition of PI3K-AKT signaling blocks PGE2-induced COX-2 expression in lung adenocarcinoma. Onco Targets Ther. 2020;13: 8197–8208. doi:10.2147/OTT.S263977

22. Xie Y, Qi Y, Zhang Y, Chen J, Wu T, Gu Y. Regulation of angiogenic factors by the PI3K/Akt pathway in A549 lung cancer cells under hypoxic conditions. Oncol Lett. 2017;13: 2909–2914. doi:10.3892/ol.2017.5811

23. Yang HL, Thiyagarajan V, Shen PC, Mathew DC, Lin KY, Liao JW, et al. Anti-EMT properties of CoQ0 attributed to PI3K/AKT/NFKB/MMP-9 signaling pathway through ROS-mediated apoptosis. Journal of Experimental and Clinical Cancer Research. 2019;38. doi:10.1186/s13046-019-1196-x

24. Ginovyan M, Petrosyan M, Trchounian A. Antimicrobial activity of some plant materials used in Armenian traditional medicine. BMC Complement Altern Med. 2017;17: 1–9. doi:10.1186/s12906-017-1573-y

25. Ginovyan M, Hovhannisyan S, Javrushyan H, Sevoyan G. Screening revealed the strong cytotoxic activity of Alchemilla smirnovii and Hypericum alpestre ethanol extracts on different cancer cell lines. 2022;10: 12–22. doi:10.3934/biophy.2023002

26. Koss-Mikolajczyk I, Kusznierewicz B, Namiesnik J, Bartoszek A. Juices from non-typical edible fruits as health-promoting acidity regulators for food industry. LWT - Food Science and Technology. 2015;64: 845–852. doi:10.1016/j.lwt.2015.06.072

27. Koss-Mikolajczyk I, Kusznierewicz B, Wiczkowski W, Sawicki T, Bartoszek A. The comparison of betalain composition and chosen biological activities for differently pigmented prickly pear (Opuntia ficus-indica) and beetroot (Beta vulgaris) varieties. Int J Food Sci Nutr. 2019;70: 442–452. doi:10.1080/09637486.2018.1529148

28. Yang X, Wang Y, Luo J, Liu S, Yang Z. Protective effects of YC-1 against glutamate induced PC12 cell apoptosis. Cell Mol Neurobiol. 2011;31: 303–311. doi:10.1007/s10571-010-9622-9

29. Seeliger D, De Groot BL. Ligand docking and binding site analysis with PyMOL and Autodock/Vina. J Comput Aided Mol Des. 2010;24: 417–422. doi:10.1007/s10822-010-9352-6

